# Intravital imaging of real-time endogenous actin dysregulation in proximal and distal tubules at the onset of severe ischemia-reperfusion injury

**DOI:** 10.1101/2021.03.01.433337

**Authors:** Peter R. Corridon, Shurooq H. Karam, Ali A. Khraibi, Anousha A. Khan, Mohamed A. Alhashmi

**Author notes:** **Corresponding Author Contact Information:** Peter R. Corridon, PhD, Assistant Professor, Department of Immunology and Physiology, College of Medicine and Health Sciences, Director, Pre-Medicine Bridge Program, College of Arts and Sciences, Khalifa University of Science and Technology, PO Box 127788, Abu Dhabi, UAE, Office. **Co-author Email Contact Information:** Shurooq H. Karam;, Ali A. Khraibi;, Anousha A. Khan;, Mohamed A. Alhashmi.

## Abstract

Severe renal ischemia-reperfusion injury (IRI) can lead to acute and chronic kidney dysfunction. Cytoskeletal modifications are among the main effects of this condition. The majority of studies that have contributed to the current understanding of IRI have relied on histological analyses using exogenous probes after the fact. Here we report the successful real-time visualization of actin cytoskeletal alterations in live proximal and distal tubules that arise at the onset of severe IRI. To achieve this, we induced fluorescent actin expression in these segments in rats with hydrodynamic gene delivery (HGD). Using intravital two-photon microscopy we then tracked and quantified endogenous actin dysregulation that occurred by subjecting these animals to 60 minutes of bilateral renal ischemia. Rapid (by 1-hour post-reperfusion) and significant (up to 50%) declines in actin content were observed. The decline in fluorescence within proximal tubules was significantly greater than that observed in distal tubules. Actin-based fluorescence was not recovered during the measurement period extending 24 hours post-reperfusion. Such injury decimated the renal architecture, in particular, actin brush borders, and hampered the reabsorptive and filtrative capacities of these tubular compartments. Thus, for the first time, we show that the combination of HGD and intravital microscopy can serve as an experimental tool to better understand how IRI modifies the cytoskeleton *in vivo* and provide an extension to current histopathological techniques.

## INTRODUCTION

Ischemia-reperfusion injury (IRI) is a complex cascade of events that support structural and functional losses in renal tubular segments. Sudden and temporary restrictions to blood flow induce oxidative stress and inflammatory responses. These responses adversely affect the vascular endothelium and tubular epithelium, hampering renal reabsorption and filtration. Depending on the severity of the IRI, this condition can be reversed to reinstate normal function or be sustained for the subsequent loss of function [1]. Severe IRI is a common cause of acute kidney injury (AKI) [2], and produces irreversible damage that supports the progression of AKI to chronic kidney disease (CKD) and, ultimately, end-stage renal failure [3]. This disease progression is a growing global health problem with no current specific treatment that has been the focus of research for several decades [3].

Animal models have been pivotal for investigating the cascade of events that support the development of irreversible damage from IRI, and have identified the proximal [4] and distal [5] tubules as major sites of injury with this condition. Severe renal IRI is characterized by losses in brush border components and polarity in proximal tubule epithelial cells, tubular occlusions, as well as cytoskeletal dysregulation [6]. Modifications in the cytoskeleton are among the main effects of IRI. However, to a lesser extent, distal segments succumb to the effects of IRI, and research has highlighted the formation of casts within the lumen of these tubules as a major manifestation of the insult. Specifically, it has been shown that cellular blebs aggregate with other intraluminal materials to form casts, which are either excreted into the urine or stay logged in the lumen and create substantial obstructions within these tubules [5]. Irrespective of the tubular segment, these changes occur rapidly and correlate with the severity and duration of insult, and affect the overall structural and functional integrity of renal tubules [7]. However, the cellular and subcellular mechanisms responsible for this cascade and resulting tubular damage, are not fully understood.

Pioneering works that have contributed to the current understanding have relied on histological analyses after the fact [8]. Furthermore, the ability to conduct these studies *in vivo* was traditionally dependent on the use of exogenous probes that provide indirect measures of live processes [9] and techniques that could only facilitate the imaging of subcellular structures in confined regions within the kidney [10]. For decades, numerous investigators have outlined the value of gene delivery to the kidney in attempts to expand the expression of trackable endogenous probes [11]. Yet, the complex nature of the renal system has provided significant challenges to progress in this field.

Recent advances in renal gene delivery have provided a much-anticipated option to address this challenge. By altering hydrodynamic fluid pressures, it has been shown that it is possible to transiently increase intravascular pressure within peritubular capillary networks and induce exogenous gene expression in the surrounding tubular epithelium [12]. This method, which is known as hydrodynamic gene delivery (HGD), is capable of producing widespread genetic alterations in various segments of the kidney, namely the proximal and distal convoluted tubules, with minimal effect to the organ [13, 14]. Therefore, to improve the current understanding of renal IRI, we extended this approach to investigate real-time changes in tubular structures that are altered in response to IRI, using a sophisticated imaging tool.

In this study, we visualized changes that occur in live proximal and distal tubules by inducing fluorescent actin protein expression in these segments in rats using HGD, subjected these animals to 60 minutes of bilateral ischemic injury, and monitored the immediate changes that occurred within the tubules after blood flow was reinstated to the kidney. Accordingly, we described and validated the combinative use of HGD and intravital two-photon microscopy as an experimental tool to track and quantify actin cytoskeletal dysregulation that occurs at the onset of severe IRI *in vivo*.

## RESULTS

### Comparative View of the Actin-Rich Brush Border *Ex Vivo* Using Exogenous and Endogenous Probes

Brightfield images were obtained from cortical sections of normal rat kidneys that were counterstained with hematoxylin and eosin (H&E) (Fig. 1A). These images outlined the intrinsic actin brush border by the localization of the exogenous eosin fluorophore (pink fluorescence), which is a hallmark of the proximal tubule that is used to differentiate it from other portions of the renal tubule in standard histological analyses. Confocal laser scanning micrographs (Fig. 1B) also revealed the actin-rich brush border using another exogenous fluorophore, Texas-red-labeled phalloidin (red fluorescence). In comparison, images collected from kidneys that received HGD to show the expression of an endogenous EGFP-actin (green fluorescence) again highlighted actin localization along the brush border in cortical sections (Fig. 1C) that correlate well with standard histological findings.

**Figure 1.**
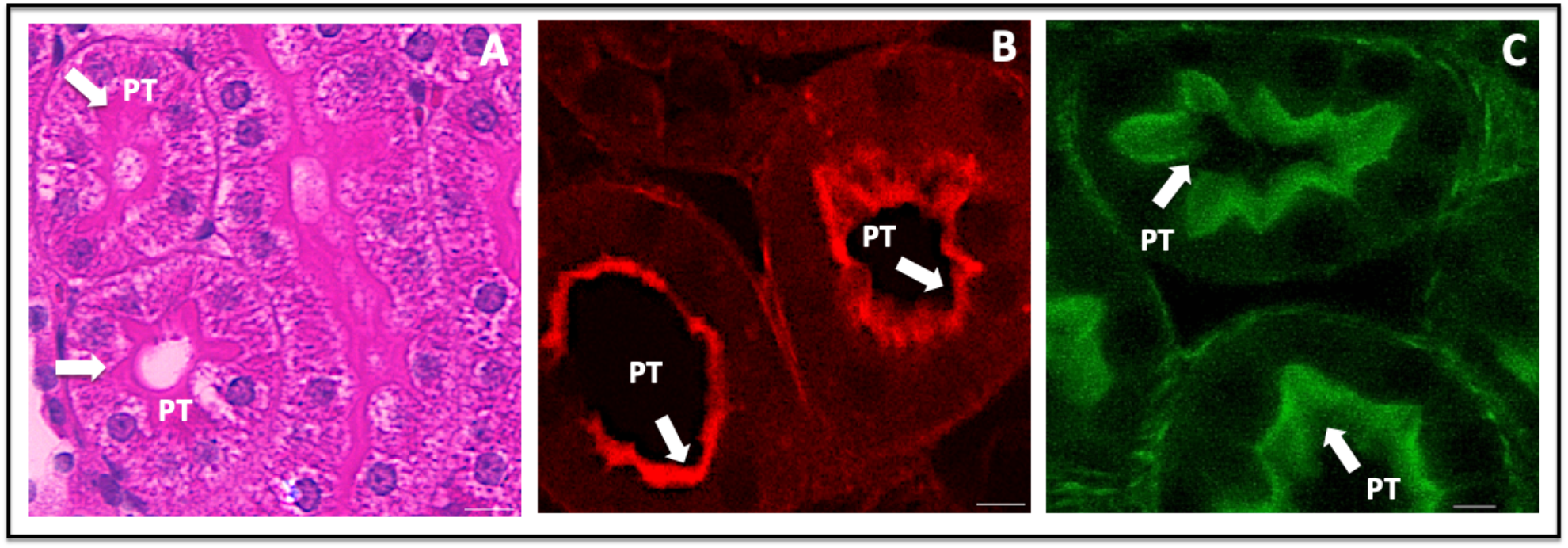
Brightfield and confocal microscopic images highlight the presence of actin in the renal brush border *ex vivo*. Images were taken with 60X objectives (2x digital zoom) using brightfield (image A) and confocal (images B and C) microscopes. These images of the proximal tubule (PT) in cortical kidney sections outline the innate actin localization (identified by arrows) along brush borders using exogenous probes (H&E, image A, and Texas-red phalloidin, image B). Similarly, image C highlights the intense presence of actin along the brush border in the PT of cortical kidney sections obtained from rats that expressed EGFP-actin fusion proteins using HGD. Image B was taken using only the red-pseudo-color channel, and image C was taken using only the green-pseudo-color channel. Scale bars represent 10 µm.

### Endogenous Fluorescent Actin Expression, and Verification of Normal Renal Morphology and Function Before Injury *In Vivo*

Fluorescent images were acquired from live kidneys and provided the ability to distinguish between proximal and distal tubules in vivo based on their relative levels of innate autofluorescence (Fig. 2A). Furthermore, an enhanced level of contrast was generated by the expression of EGFP-actin, primarily along the brush border (Fig. 2B through Fig. 2E) and correlate with the distribution of actin observed using conventional histological techniques (Fig. 1). Such fluorescent protein expression provided an additional way to distinguish between proximal and distal segments, which are routinely visualized *in vivo* using this imaging technique and allowed us to monitor the actin cytoskeleton. Moreover, continuous imaging over a period of 60 minutes did not appear to induce photobleaching (Fig. 3).

**Figure 2.**
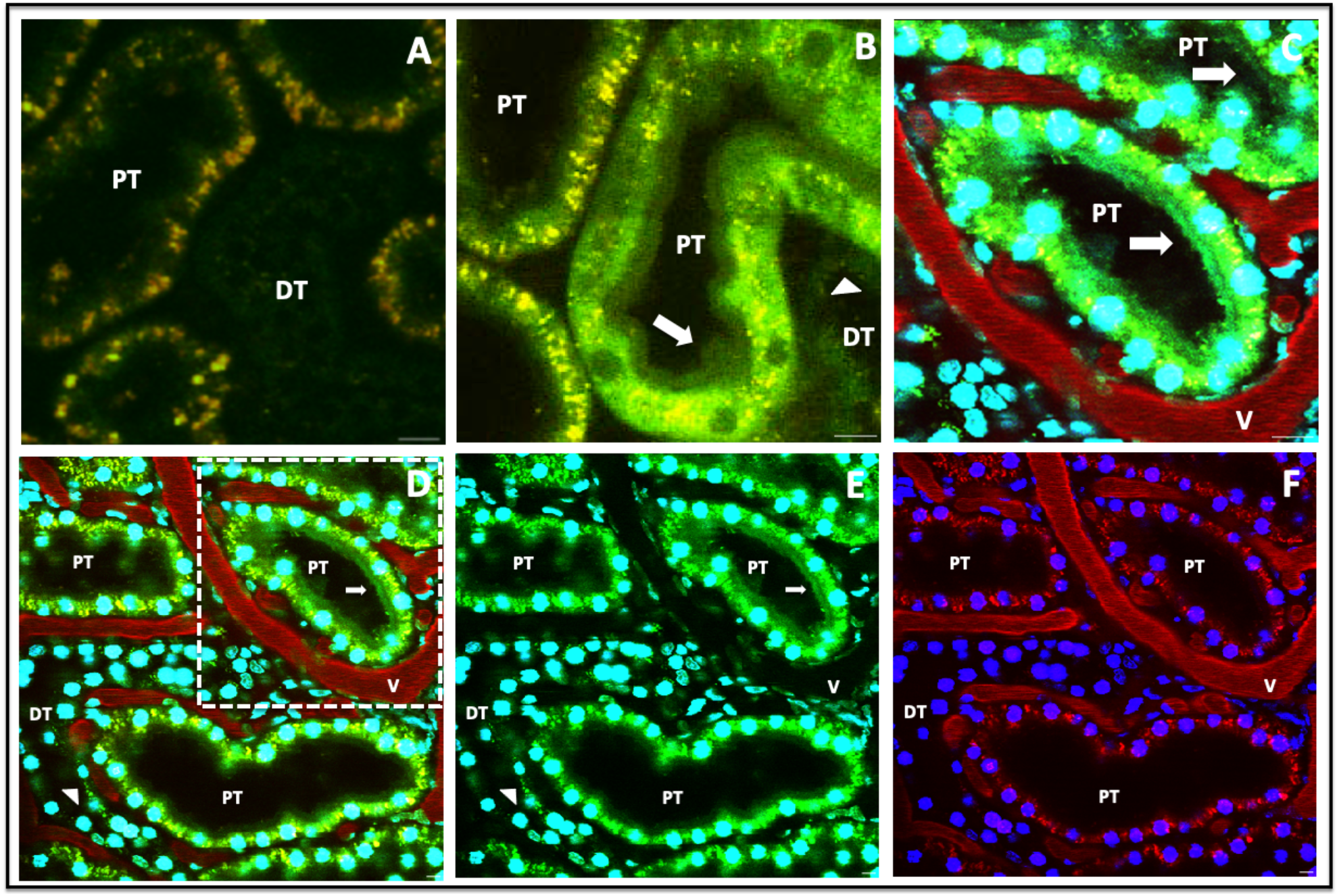
HGD allowed the visualization of the actin-rich renal brush border *in vivo*. Image A (taken at 2X optical zoom) shows innate autofluorescent patterns that are used to routinely distinguish the proximal tubule (PT) from the distal tubule (DT) segments but were unable to outline brush border segments. In comparison, images B and C (taken at 2X optical zoom), as well as D and E (taken at 1X optical zoom), highlight the presence of the actin-rich brush border *in vivo* (identified by arrows) in the proximal tubule. The region outlined in image D (dashed-line) is presented as image C, to focus on the brush border as we did in Fig. B. Images A and B were formed by merging the green- and red-pseudo-color channels, while images C and D were formed by merging the blue-, green- and red-pseudo-color channels. We presented different combinations of the pseudo-channels shown in image D to create images E and F, to better highlight EGFP-actin expression in the tubules. Specifically, Image E was created by merging the green- and blue-pseudo-colors, and image F was created by merging the blue- and red-pseudo-colors. Overall, the presence of Hoechst 33342 and 150-kDa TRITC-dextran dyes in images C through F delineated the tubular and supporting vasculature architectures. Scale bars represent 20 µm.

**Figure 3.**
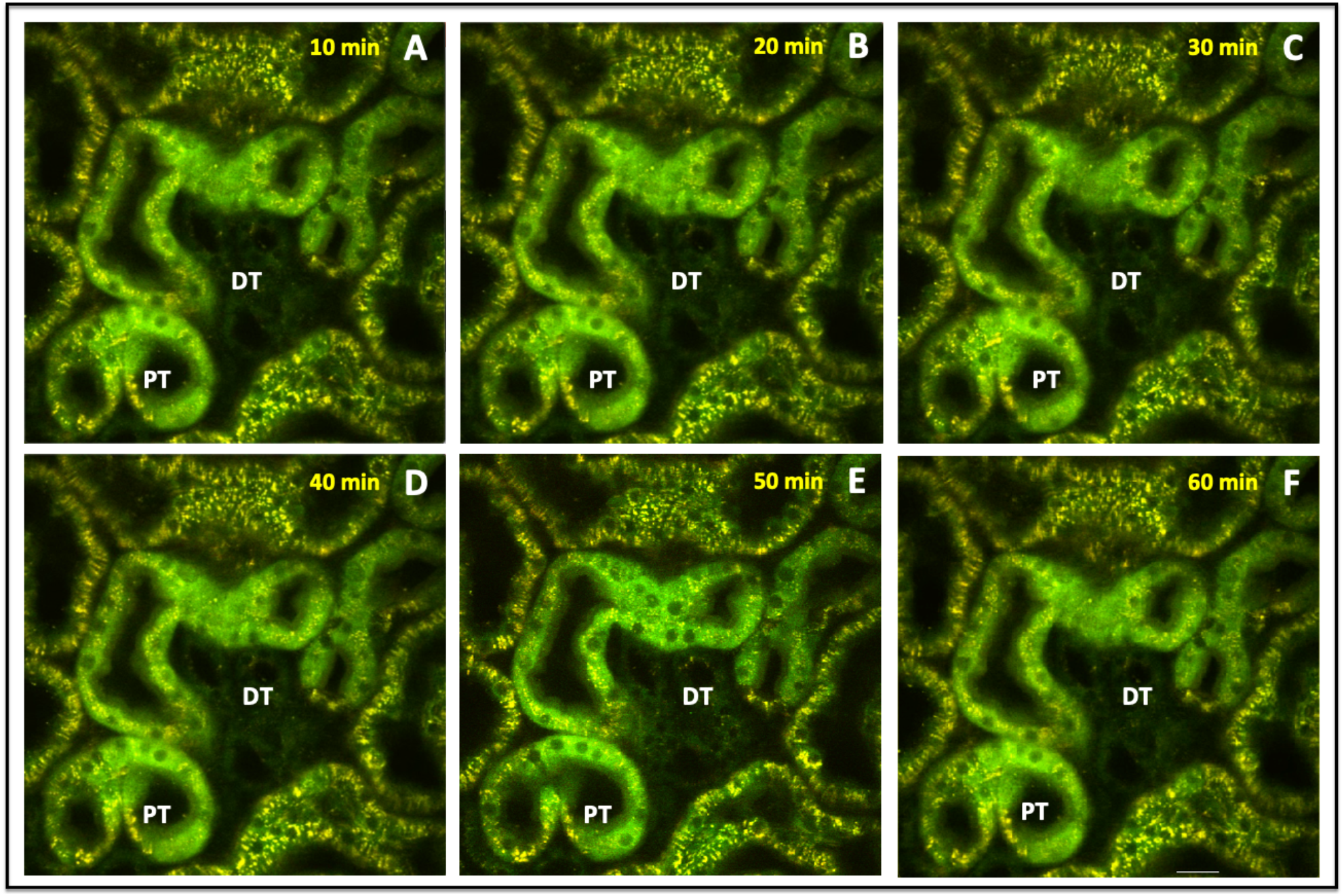
Intravital two-photon micrographs were taken with a X60 objective from a live rat that received HGD. All images were formed by merging the green-and red-pseudo-color channels and shows the homogenous distribution of EGFP-actin expression in both proximal and distal tubules that did not appear to be affected by continuous imaging over a 1-hour period. Scale bar represents 20 µm.

EGFP-actin expression also outlined standard morphology that would support innate functions, such as patent lumens of proximal and distal tubules. The venous introduction of Hoechst 33342 and 150-kDa TRITC-dextran dyes also supported the histological verification of inherent functional morphology in the rat kidney before IRI (Fig. 2C and 2F). Hoechst 33342 stained the nuclei of proximal tubular epithelial cells to display their typical appearance, and the highmolecular-weight dextran molecules were confined to the lumen of peritubular capillaries and confirmed normal vascular architecture.

### Impact on *In Vivo* Tubular Structure and Function with Severe Ischemia-Reperfusion Injury

This form of injury decimated the renal architecture, in particular, the actin brush border, and hampered the reabsorptive and filtrative capacities of these tubular compartments. After 1 hour of reperfusion, live imaging provided evidence of alterations to normal tubular structure and function (Fig. 4B and 4D). The disruptions to tubular EGFP-actin fluorescence, as well as innate autofluorescence, made it difficult to distinguish proximal from distal tubular segments.

**Figure 4.**
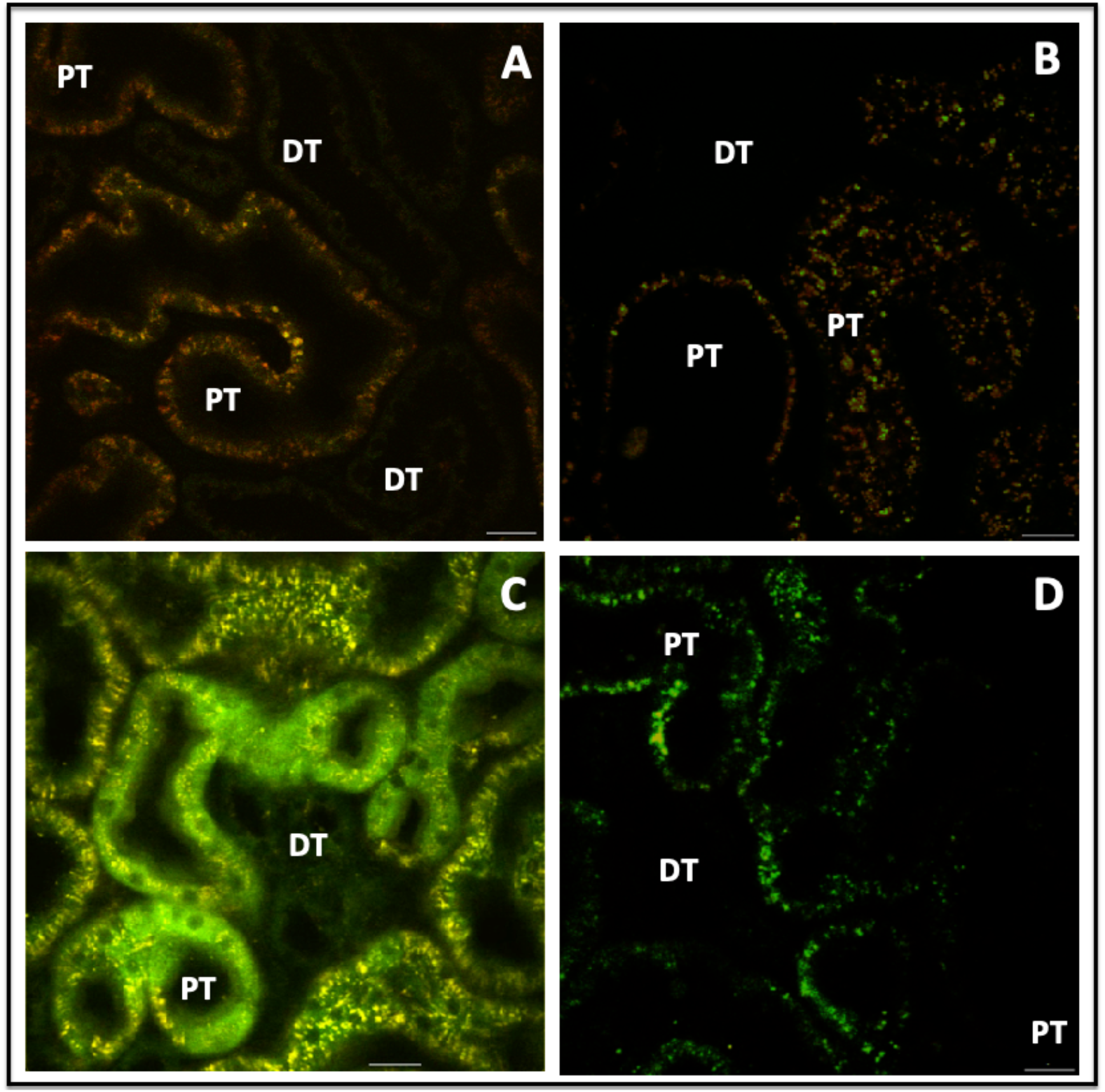
Extensive alterations to tubular structure that occurred after one hour of reperfusion. Intravital two-photon micrographs taken with a 60X objective show the effects of severe IRI *in vivo*. Animals that received sham injuries maintained intact tubular structure (images A and C). Image A was obtained from an animal that did not receive HGD (group 1), while image C (which displays the actin-rich brush border) was obtained from an animal that received HGD (group 3). In comparison, we also observed substantial damage to both proximal and distal tubules in images B and D, which were taken from animals that were subjected to severe IRI (animals in group 2 did not receive HDG, image B, and animals in group 4 received HGD, image D). Images C and D can also be found in Fig. 2 (as image F) and Fig. 6 (as image F) respectively. Such injury dysregulated the actin cytoskeleton, and specifically, stripped the proximal tubule of its characteristic brush border that was visible in (image C). Scale bars represent 20 µm.

Moreover, after 24 hours of reperfusion, the introduction of fluorescently labeled low-molecular-weight (4-kDa FITC) and high-molecular-weight (150-kDa TRITC) dextran markers provided further evidence of distorted renal function (Fig. 5). For instance, the combined presence of the FITC and TRITC dextrans within the lumen of the tubules outlined that both types of molecules could have been simultaneously filtered by glomeruli. The combined presence of these dyes within the lumen highlighted the possible impairment of normal filtrative capacities, as the molecular-weight should have been confined to the vasculature, and not enter the filtrate. Imaging of these regions over a subsequent period of 1 hour confirmed the severity of the induced renal injury (Supplemental Video 1). We also observed aggregated red blood cells (rouleaux) within vasculature, sluggish blood flow and narrowed peritubular capillaries. Analogously, there was little evidence to support the entry of low-molecular-weight dextran molecules within the proximal tubules. This process may indicate the impairment of innate tubular endocytic capacities and correlate with the dysregulation of the actin cytoskeleton and induced injury.

**Figure 5.**
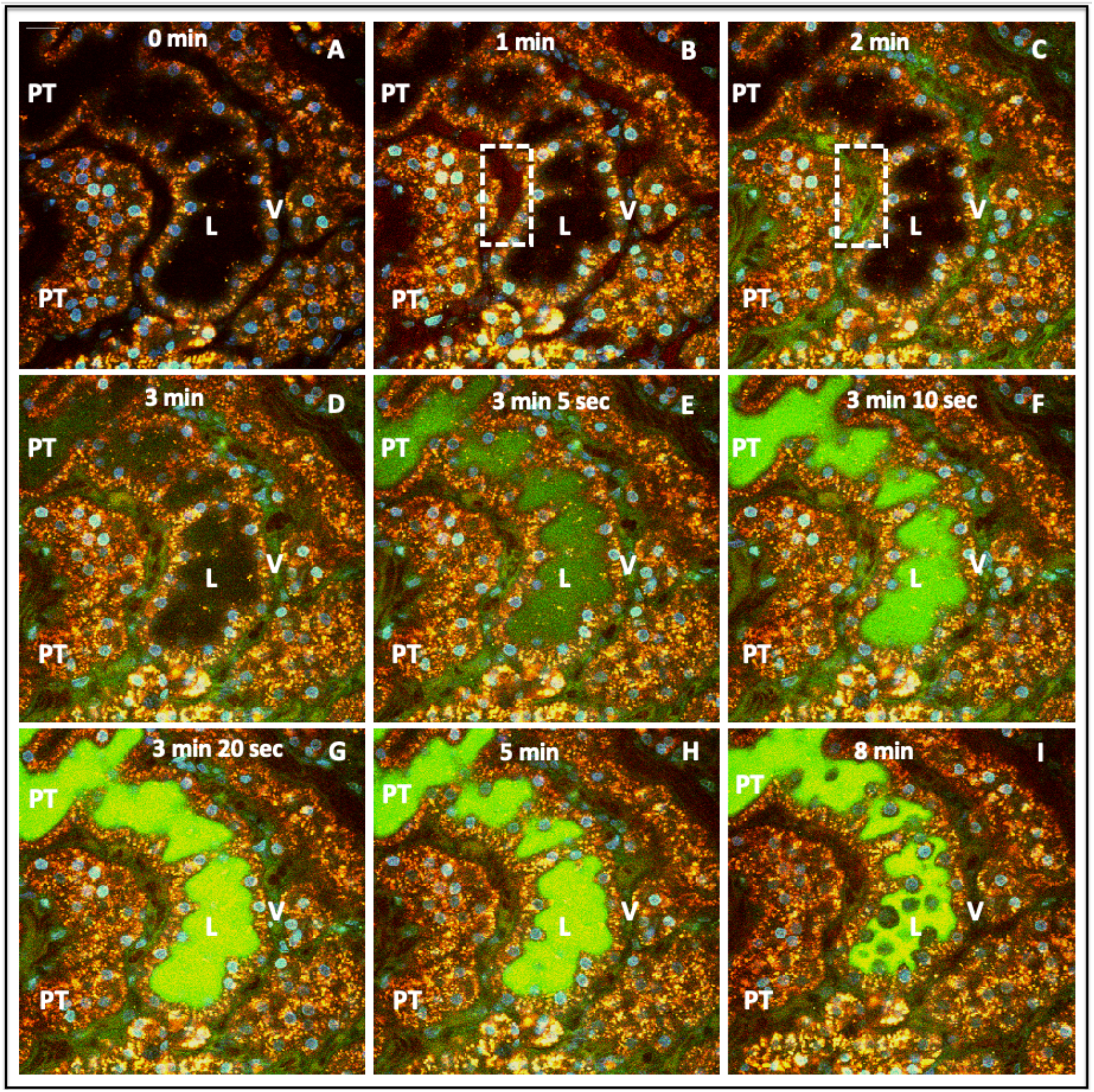
Time-lapse images outline disruptions to normal renal filtrative and endocytic capacities that resulted from severe IRI. Intravital two-photon micrographs taken with a 60X objective from a live rat in group 4, which received HGD and was subjected to IRI. After reinstating blood flow to the kidney, we observed a substantial injury 24 hours after reperfusion. At that time point, we infused of a mixture of 4-kDa FITC and 150-kDa TRITC dextrans, along with Hoechst 33342, via the jugular vein of the animal, to track renal dynamics. Images A through I illustrate the loss of EGFP-actin expression, reductions in the thicknesses of the vasculature (V), aggregated red blood cells (rouleaux) in the peritubular capillaries (dashed line in image D), absence of endocytic uptake of low-molecular-weight FITC dextran molecules by the proximal tubules, and simultaneous entry of both FITC and TRITC dyes in the lumen. Moreover, these images illustrate the initial presence of the TRITC dye entering the peritubular vasculature (image B) and then the entry of the FITC dye (image C). After that, there was a reduced level of fluorescence within the vasculature observed in image I. Scale bar represents 20 µm. The time-lapse video for this event is presented in Supplemental Video 1.

### Real-Time *In Vivo* Imaging of Actin Dysregulation at the Onset of Reperfusion

We first examined the changes in fluorescence intensity within individual groups across the first 60 minutes of reperfusion in the sham injury and IRI models. There were significant differences in fluorescence intensity between proximal and distal tubular segments in rats in group 1 (no gene delivery, sham injury), based on native autofluorescence (p = 0.015), and those in group 4 (gene delivery, IRI), based on EGFP-actin fluorescence (p=0.023). Whereas analyses performed on animals in groups 2 (no gene delivery, IRI) and 3 (gene delivery, sham injury) showed that the analogous reductions in fluorescence were not significant, (p = 0.078) and (p = 0.428), respectively.

Using data recorded from the two groups of animals that received sham injuries (group 1 and group 3), the Student’s t-test identified a significant difference (p = 0.006) in the reductions of proximal tubular fluorescence intensity, but not in the decreases in distal tubular fluorescence intensity (p = 0.237), between these groups. In comparison, data obtained from animals subjected to IRI (group 2 and group 4) revealed significant differences between the loss in fluorescence intensity in proximal tubules in group 2 and those in group 4 (p = 0.004). We also observed significant differences between the loss in fluorescence intensity in distal tubules in group 2 and those in group 4 (p = 0.003). Additionally, the decline in actin-based fluorescence intensity in proximal tubules was significantly greater than that observed in distal tubules among rats in group 4 (p = 0.027). We also compared data among the four groups and assessed the effect of injury using the ANOVA test. The observed F value 14.436 is larger than the critical value of 3.098 and may be interpreted as statistically significant difference among the means of the groups at the α error level 0.05 (p = 3.029 × 10^−5^).

Fluorescent actin protein expression allowed us to characterize general, as well as specific actin-based, alterations that were observed in rat proximal and distal tubules (Fig. 6). Using the data, we quantified the time-dependent variations in fluorescence intensity (Fig. 7). Imaging was conducted in a manner previously utilized to limit the occurrence of phototoxicity [15], and was confirmed in control studies presented in Fig. 3. Within 10 minutes of reperfusion, tubular lumens were narrowed, and normal actin-rich brush border patterns were replaced by coalesced masses that blocked the lumen of 30-40% of the tubules that were imaged. EGFP-actin appeared more heterogeneously distributed and clumped at that time, yet it was still possible to differentiate between distal and proximal tubules then (Fig. 6A).

**Figure 6.**
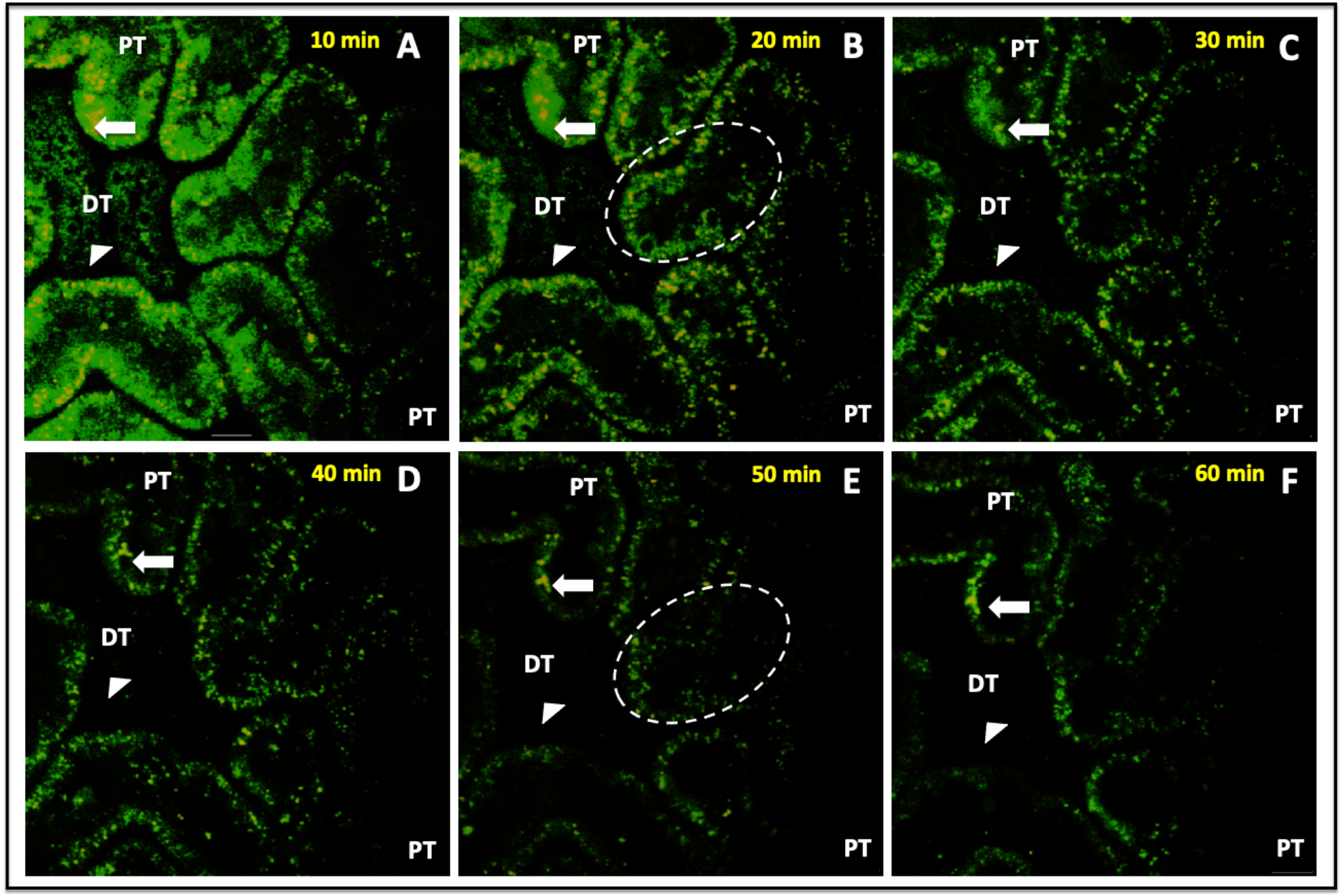
Time-lapse images tracked alterations in actin-based fluorescence observed during the first 60 minutes of reperfusion. Intravital two-photon micrographs taken with a 60X objective from a live rat in group 4 across 60 minutes (this is the same imaging field that is previously presented in Fig. 3D). This animal received HGD and was subjected to ischemia-reperfusion injury (IRI). All images were formed by merging the green-and red-pseudo-color channels and shows the expression of EGFP actin in both proximal and distal tubules. This fluorescent protein expression allowed us to visualize the live and real-time changes in tubular structure and function that resulted from IRI (arrows identified changes in proximal tubules, and arrowheads identified changes in distal tubules). At the 10-minute mark, EGFP-actin appeared more heterogeneously distributed and clumped in tubular segments. The dashed ovals in images B and E track the outlined region and show how cells have sloughed off the proximal tubule segment and migrated into the lumen to generate ghost tubules (tubules mostly devoid of living cells) by the 50-minute mark. It should be noted that there were minor shifts in the field during the 60-minute imaging period that resulted from the vibration caused by respiration. Scale bar represents 20 µm. A time-lapse video showing portions of this event is presented in Supplemental Video 2.

**Figure 7.**
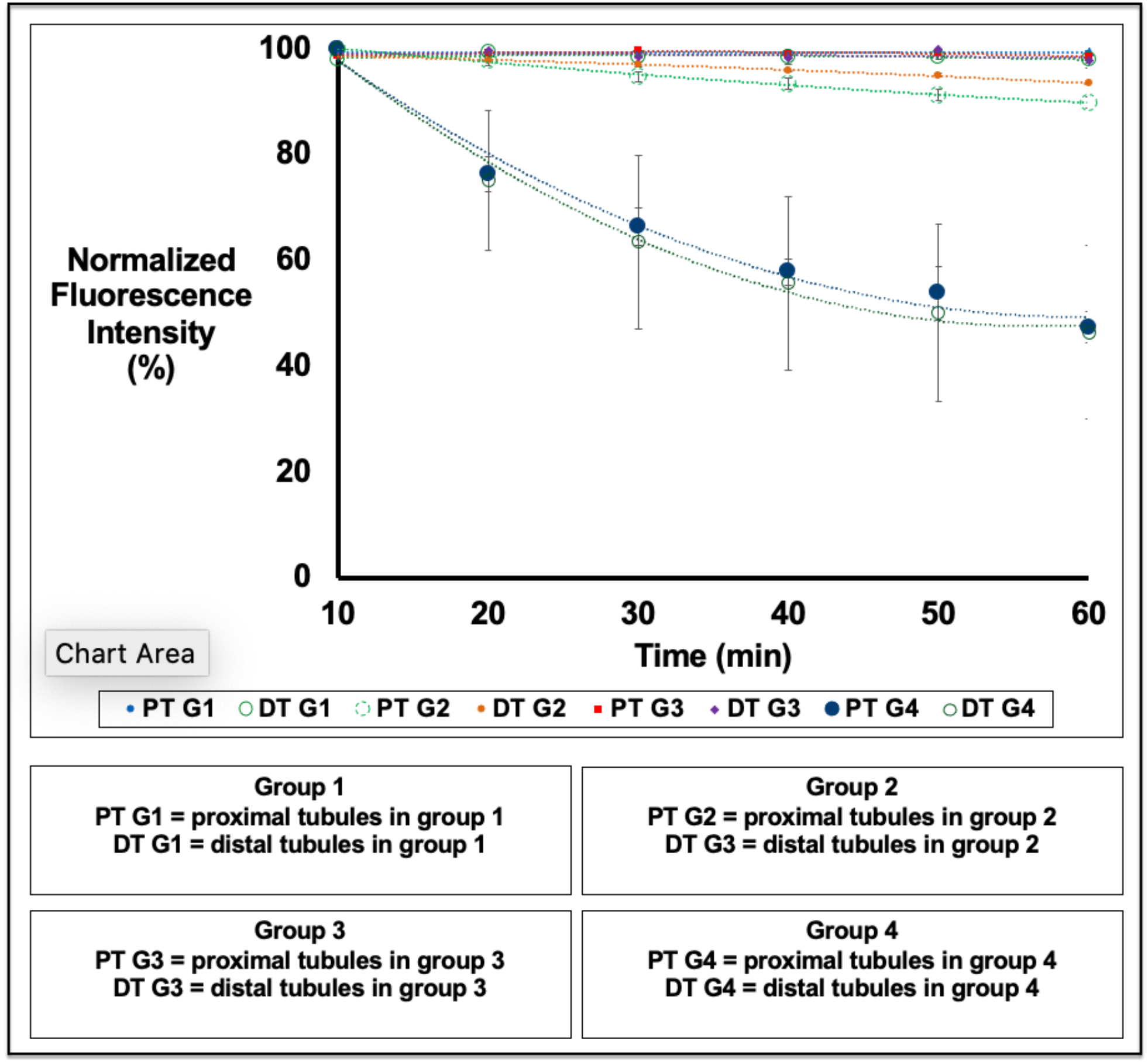
*In vivo* changes in mean fluorescence intensities obtained from proximal and distal tubular segments. There were no considerable differences in fluorescence intensity recorded from proximal and distal tubular segments from animals that did not receive HGD (group 1), but there were larger variations in autofluorescence that resulted from ischemia-reperfusion injury (group 2) across the 60-minute measurement period. In comparison, we observed substantial decreases in fluorescence intensities in proximal and distal tubular segments recorded from animals that received HGD (groups 3 and 4).

As time progressed, fluorescent clumps were dislodged from the tubule into the lumen. This sloughing process continued, and fluorescent actin-derived structures amalgamated into free-floating blebs and casts of various sizes within the lumen (Fig. 6B and 6C). Antegrade flow within the tubules supported the movement of the fluorescent cell/tissue debris through the lumen (Supplemental Video 2). This normal flow pattern was intermittently replaced by retrograde flow that accompanied further abnormal tubular narrowing.

By the 20-minute mark (Fig. 6B), there was an intense, approximately 30%, reduction of EGFP-actin fluorescence in both proximal and distal tubular components. At that time, it was difficult to find signs of intact brush borders, as the majority of these components were shed into the lumina leaving behind greater heterogeneity in fluorescent actin localization. Moreover, it was possible to witness entire groups of cells, within the cuboidal epithelia, dislocate from their tubular linings. Fluorescent debris was restricted to the lumen, as there were no signs of EGFP-actin in regions that corresponded to the neighboring vasculature. These changes supported the development of ghost tubules (tubular segments devoid of living cells, previously identified by Hall et al. using intravital multiphoton microscopy [16]), and, in some instances, we observed drastic improvements in the patency of the tubular lumen, as fluorescent debris was seen to be transported swiftly and bidirectionally within the lumen (Supplemental Video 2).

Furthermore, after 40-minutes of reperfusion (Fig. 6D), we observed the decimation of 20-30% of all imaged tubular segments, and it became difficult to locate and even dentified the lineage of various tubules after 50 minutes of IRI (Fig. 6B). The progressive loss of fluorescence continued, and resulted in substantial declines in EGFP-actin fluorescence within the first hour of reperfusion (Fig. 6A through 6F). Finally, we estimated as much as 60% reductions in EGFP-actin fluorescence occurred in both proximal and distal segments after 60 minutes reperfusion. Interestingly, actin-based fluorescence was not recovered during our measurement period that extended to roughly 24 hours after reperfusion.

## DISCUSSION

Fluorescent probes and animal models have been used extensively to investigate mechanisms that generate irreversible damage from IRI. Gaining a better understanding of the disease etiology can help devise novel strategies to prevent the progression of AKI to CKD and, ultimately, kidney failure. Traditional light and electron microscopy have provided significant insight into the cascade of events that occur with such pathologies [12]. Yet, consolidated and unified descriptions of the associated cellular and sub-cellular mechanisms are needed. A major technical drawback that has limited progress in this area lies in the ability to perform these investigations *in vivo*.

Recent advances in imaging technologies and genetic engineering have provided a means to perform such studies in real-time. Powerful imaging tools, like intravital two-photon microscopy, have contributed to the present understanding of the functional morphology in the live kidney, deviations that occur with damage, and ways to better manage these conditions [16]. Likewise, techniques like HGD can help change the genetic makeup of cells within the kidney, and thus offer a newfound way to examine *in vivo* processes using endogenous markers [17]. As a result, in this study, for the first time we show that the combination of intravital two-photon microscopy and HGD can be used to visualize and measure the rate of degradation of the actin cytoskeleton at location known to be targeted by IRI.

Our *ex vivo* findings illustrate that HGD can facilitate the expression of genetically altered forms of actin within proximal and distal tubules. The fluorescence patterns collected from cortical kidney sections, using confocal microscopy, confirm the intrinsic localization of EGFP-actin fusion proteins, particularly along the brush border. This is an important histological characteristic that has been relied on for decades [8], and that has been previously recorded *ex vivo* using micropuncture gene delivery [9].

In comparison, for *in vivo* imaging studies, the early proximal and distal tubules in rodents can be easily accessed for live imaging using intravital two-photon microscopy, and routine distinctions are made between these tubules based on their relative levels of autofluorescence [18]. Furthermore, the fluorescent images acquired from live kidneys highlighted the different renal structural patterns observed *in vivo* and *ex vivo* as previously reported, while confirming the enhanced presence of actin along the brush border [19]. Proximal segments have higher autofluorescent signatures than their distal counterparts. However, it is difficult to differentiate between the individual segments of the proximal convoluted tubule based only on innate tissue autofluorescence. We thus sought to determine whether HGD could provide a better means for tubular segment differentiation. We observed that hydrodynamic-based renal gene transfer facilitated the endogenous expression of actin fusion proteins within the renal tubules, and thus enhanced the contrast between distal segments and proximal tubules.

Overall, EGFP-actin expression helped outline normal renal morphology and function along with the nuclear and vascular probes in live rodent kidneys. To further underscore the utility of the model, live EGFP-actin expression was visualized within and along the brush border of proximal tubular epithelial cells at various levels, and thus the varied degrees of actin localization can provide a means to distinguish between S1 and S2 segments of the proximal tubule, based on the relative thicknesses of the actin brush border, similar to pioneering studies conducted that provided this distinction in cortical segments [8]. Our studies used plasmid transgene vectors that expressed both fluorescent filamentous (F-actin) and monomeric globular (G-actin) proteins, and thus additional segment-specific markers will be needed to support this claim. In the future, we can also consider the use of plasmid vectors that support the fluorescent expression of only F-actin fusion proteins [20] to refine the way actin cytoskeletal components can be tracked *in vivo*. However, these investigations may be limited by the resolution of the imaging system.

Also, in some instances there was a higher fluorescent signal from non-filamentous actin, which is consistent with previous research conducted in cell culture [20]. These *ex vivo* studies have also determined that the expression of eGFP-actin can affect cell behaviour. Fortunately, this concern as the plasmid titer and period of expression were previously shown to support the stable expression of fluorescent/exogenous proteins that did not significantly alter cellular function *in vivo* [13, 14]. It was suitable to consider this type of expression vector for our initial studies, as both forms of actin are essential cytoskeletal components.

Once the focus was shifted to investigating the impact of injury, it was evident that this severe form of IRI had a devastating effect on tubular structure and function. Bilateral renal ischemia for the 60-minute period would have supported severe and sustained reductions in blood pressure to induce tubular necrosis [21]. Pathological changes that occur with this condition include reduced filtrative capacities of the tubules that results from hypoperfusion. Cellular debris and casts also amalgamate to obstruct the lumina and hinder the movement of the filtrate through the nephron.

The damage to the tubular epithelium that stems from ischemia has conventionally been considered as a consequence of cellular necrosis [22], and these irreversible effects, which were visualized in real-time within the first 60 minutes of our reperfusion study, demonstrate the proof of concept. However, there is growing evidence to suggest that apoptosis has a significant contribution to the acute injury. Reductions in renal apoptosis antagonizing transcription factor have been shown to result from IRI, hampering intrinsic activation of antiapoptotic pathways and/or inhibition of proapoptotic pathways [23]. Further investigations that can combine this imaging technique and gene delivery may be used to identify the potential therapeutic application of this transcription factor in IRI, and potentially extend the value of the presented model.

Meanwhile, it is well known that the proximal tubules rely mainly on mitochondrial metabolism for ATP synthesis due to their limited glycolytic capacities and are thus particularly susceptible to IRI. The rapid and significant recorded reductions in actin-based fluorescence allowed us to visualize such proximal tubular damage, which included brush border losses that would have resulted from profound decreases in intracellular ATP. Drops in ATP levels would have occurred early after onset of ischemia and driven actin cytoskeletal derangements that favor the non-filamentous form of actin, as the cytoskeleton requires ATP to remain in a filamentous form [16].

Cytoskeletal dysregulation would have, in turn, led to the redistribution of integrins and Na^+^-K^+^-ATPase from the basal membrane [24]. This process would have resulted in impaired cellular transport mechanisms that ultimately support cellular death and sloughing from tubular basement membranes. Comparatively, the distal tubule epithelium would have succumbed to less damage based on their relatively lower dependence on mitochondrial metabolism but would have been drastically impacted by intraluminal obstructions generated from damage to proximal tubular segments [5, 21]. Paradoxically, reinstating blood flow would have supported additional tissue damage. Emerging evidence suggests that the mitochondrial production of reactive oxygen species, which occurs during the reperfusion phase of IRI, has a critical role in destroying cellular components, as well as initiating apoptosis and necroptosis [25]. Monitoring this process thus allowed us to quantify the cumulative damage by estimating the reductions in EGFP-actin fluorescence that occurred within the first hour of reperfusion. Furthermore, the ability to examine live events with this novel approach, like antegrade flow patterns that occurred within the tubular lumen, as well as the dynamic changes in tubular diameter extend beyond the limits of established histopathological techniques.

In summary, due to the complex nature of the kidney, *in vivo* studies have relied on exogenous probes to investigate the underlying nature of renal tubular morphological and functional processes [26, 27]. To extend this approach, this study demonstrates the utility of the combinative use of HGD and intravital two-photon microscopy to track the dynamic remodeling of the actin cytoskeleton. Importantly, this method signifies a way to monitor intrinsic cellular and molecular mechanisms involved in the generation of irreversible kidney injury that results from IRI. Future studies can be employed to compare the levels of alterations in and potential recovery of actin content as a function of injury severity, tubular complexity and renal function.

Furthermore, such studies can reinforce the combined use of HGD and intravital imaging. This combination may provide a powerful tool to examine therapeutic targets that can limit the progression of renal injuries associated with IRI [28].

## METHODS

### Fluorescent Plasmids and Dyes

Plasmid DNA encoding enhanced green fluorescent (EGFP)-actin (Takara Bio USA, Mountain View, CA) facilitated exogenous gene expression in rodent kidneys. The following dyes were used for intravital two-photon imaging and bolus injected intravenously in a volume of 0.5 ml: 50 µl of 150-kDa tetramethyl rhodamine isothiocyanate (TRITC) and/or 4-kDa fluorescein isothiocyanate (FITC) dextrans (TdB Consultancy, Uppsala, Sweden) and 30-50 µl of Hoechst 33342 (Invitrogen, Carlsbad, CA). Texas red-phalloidin (Invitrogen Corporation, Mountain View, CA) was used for *ex vivo* actin staining.

### Hydrodynamic Gene Delivery

All experiments were perfromed on 200 to 400 g male Sprague-Dawley rats (Harlan Laboratories, Indianapolis, IN). The experimental were approved by the Indiana University School of Medicine Institutional Animal Care and Use Committee, and Animal Research Oversight Committee at Khalifa University of Science and Technology, and the study was carried out in compliance with the ARRIVE guidelines. Animals were anesthetized with inhaled isoflurane (5% in oxygen, Webster Veterinary Supply, Devens, MA) and then given intraperitoneal injections of 50 mg/kg of pentobarbital (Hospira, Inc., Lake Forest, IL). The details of the HGD process are outlined in the literature [13, 14]. Briefly, for this process, 1-3 µg of EGFP-actin plasmid DNA, per gram of body weight were suspended in 0.5 ml of saline for retrograde renal vein injections. Animals were allowed 14 days to recover before further experimentation.

### Brightfield Imaging

Kidneys were fixed with 4% paraformaldehyde for 24 hours at 4°C, and immersed in 4% phosphate-buffered formalin, again for a minimum of 24 hours at room temperature. Specimens were rinsed in distilled H_2_O and stored in 70% ethanol. Specimens were dehydrated through a graded series of ethanol (70%; 80%, 95%, 100%), cleared in xylene, infiltrated with 4 changes of paraffin (under vacuum at 59°C; 45 minutes each), and embedded in fresh paraffin. After which, 4-5 μm thick sections were collected with a Reichert-Jung 820 microtome (Depew, NY), flattened on a warm water bath and mounted on glass slides, and stained with H&E. A Nikon Microphot SA Upright Microscope with a 60X objective and sensitive Diagnostic Instruments SPOT RT Slider color camera (Nikon, Tokyo, Japan) used to collect images.

### Confocal Imaging

Whole kidneys were harvested from control rats and those that received HGD. Kidneys were immersion fixed with 4% paraformaldehyde, 100-200 μm thick cortical sections were obtained, and incubated overnight in a phalloidin staining solution. This solution was prepared by diluting Texas-red-phalloidin in a blocking buffer (2% bovine serum albumin and 0.1% Triton X-100, diluted in phosphate-buffered saline) at a ratio of 1:200 for roughly 24 hours. The tissues then were rinsed three times for two hours in PBS and mounted onto slides. Images were collected with a 60X objective.

### Intravital Two-Photon Imaging

While under sedation, vertical flank incisions were made to externalize left kidneys for imaging.^10^ In some cases, the internal jugular vein was cannulated for intravenous infusions of dyes. Body temperature was controlled, as exteriorized kidneys were positioned inside a glass-bottom dish containing saline, which was set above a 60X water-immersion objective. Fluorescent micrographs were collected using an Olympus (Center Valley, PA) FV 1000-MPE Microscope equipped with a Spectra-Physics (Santa Clara, CA) MaiTai Deep See laser, with dispersion compensation for two-photon microscopy, tuned to 770-860 nm excitation wavelengths. The system was mounted on an Olympus IX81 inverted microscope, was also equipped with dichroic mirrors to collect blue, green, and red emissions and two external detectors for two-photon imaging.

### Bilateral Renal Ischemia-Reperfusion Injury

Two weeks after recovering from the HGD, transfected animals, along with others that did not receive gene transfer, were separated into four groups (n=3 for all groups). Groups 1 and 2 did not receive gene transfer, while groups 3 and 4 received HGD. All animals were anesthetized for median laparotomies that allowed blunt dissection of renal pedicles. For rats in groups 2 and 4, non-traumatic vascular clamps were applied to bilateral renal pedicles simultaneously for 60?minutes. After the clamps were removed, reperfusion was confirmed visually. Whereas rats in groups 1 and 3 received sham injuries. Midline incisions were closed, and the animals were prepared for intravital imaging.

### Investigation of Changes in Fluorescence and Tubular Function

Two-photon fluorescent micrographs were collected to analyze immediate structural and functional changes in live kidneys directly after reperfusion. For morphological changes, we estimated relative variations in autofluorescence or EGFP-actin fluorescence in proximal and distal tubular segments during the first 60 minutes of reperfusion. Four equal and adjacent regions were randomly chosen on proximal and distal tubular segments to record changes in mean fluorescence intensities at 10-minute intervals. The data was averaged across each group to track time-based losses in actin fluorescence post reperfusion. Changes in tubular reabsorptive and filtrative capacities were also analyzed at the 60-minute mark using fluorescent dextrans [10].

### Statistical Analysis of Data

Statistical data are presented as the mean ± SE. Differences in fluorescence intensities were investigated among study groups using one-way analysis of variance (ANOVA) and Student t-tests were applied with p < 0.05 level of significance as appropriate.

## ACKNOWLEDGMENTS

The authors would like to acknowledge George J. Rhodes, MD (Indiana University) for contributions and guidance in developing the live injury model.

## AUTHOR CONTRIBUTIONS

Author contributions: P.R.C. conceived and designed research; P.R.C. performed experiments; P.R.C. and S.H.K. analyzed data; P.R.C. interpreted results of experiments; P.R.C., S.H.K., A.A.K., A.A.K., and M.A.A. prepared figures; P.R.C., S.H.K., A.A.K., A.A.K., and M.A.A. drafted the manuscript; P.R.C., S.H.K, A.A.K., A.A.K., and M.A.A. edited and revised manuscript; and P.R.C., S.H.K., A.A.K., A.A.K., and M.A.A. approved final version of manuscript.

## SUPPLEMENTARY MATERIAL

Video 1. Renal tubular filtrative and endocytic capacities impaired by severe ischemia-reperfusion injury. Supplemental Video 1 available at URL: https://figshare.com/s/9b9624b41b2ef7c0c751 DOI: 10.6084/m9.figshare.13615889

Video 2. Actin dysregulation at the onset of severe ischemia-reperfusion injury. Supplemental Video 2 available at URL: https://figshare.com/s/02d7148b26446224c0e3 DOI: 10.6084/m9.figshare.14130146

